# *Arabidopsis thaliana ACTIN DEPOLYMERIZING FACTOR*s play a role in leaf senescence regulation

**DOI:** 10.1101/2024.05.14.594232

**Authors:** Tomoko Matsumoto, Koichi Kobayashi, Noriko Inada

## Abstract

ACTIN DEPOLYMERIZING FACTORs (ADFs) regulate the organization and dynamics of actin microfilaments (AFs) by cleavage and depolymerization of AFs. The *Arabidopsis thaliana* genome encodes 11 *ADF* genes grouped into four subclasses. Subclass I *ADF*s, *ADF1*, *-2*, *-3*, and *-4*, are expressed in all vegetative tissues and are reportedly involved in the regulation of plant growth, and biotic and abiotic stress responses. Furthermore, the nuclear localization of ADF4 is seemingly important in disease responses. Here, we present data that indicate a previously unknown regulatory role of subclass I ADFs in leaf senescence. *ADF4* knockout mutants (*adf4*) and transgenic lines in which the expression of all members of subclass I *ADF*s was downregulated (*ADF1-4Ri*) showed acceleration of both dark-induced and age-dependent leaf senescence. Among the eight *ACTIN* genes encoded in *A. thaliana*, *ACT2*, *-7*, and *-8* are expressed in vegetative tissues. In contrast to *adf4* and *ADF1-4Ri*, neither *ACT2* and *ACT8* double mutant (*act2/8*), nor *ACT7* knockout mutant (*act7*), showed accelerated leaf senescence. Upregulation of the expression of senescence-associated genes occurred earlier in *adf4* and *ADF1-4Ri* lines than in wild type. Examination of the expression of subclass I *ADF*s genes during senescence revealed a reduced expression of *ADF4* but not of other subclass I members. Additionally, we showed that nuclear localization of ADF4 was important for regulating leaf senescence. Altogether, our data indicate that subclass I ADFs, particularly ADF4, play an important role in the regulation of leaf senescence.

## INTRODUCTION

Leaves are the primary organs of biomass production via photosynthesis. After ending their role as photosynthetic organs, leaves degrade chlorophyll and proteins, resulting in gradual yellowing of their color (Lim et al. 2007). During this process, seed development is nourished through the recycling of degraded proteins and other cellular components in the reproductive organs (Lim et al. 2007). Such degradation and recycling are part of the process of leaf senescence and is regulated by external stimuli, such as light, temperature, and ambient humidity (Bohnert et al. 1995, Quirino et al. 1999, Balazadeh et al. 2010, Tan et al. 2023). Because the absence of light precludes photosynthesis and induces this recycling process, dark treatment is a useful method to force leaf senescence for research purposes (Weaver and Amasino 2001, Zhang et al. 2018, Eckstein et al. 2021).

Both forward and reverse genetic analyses have shown that leaf senescence is genetically regulated (Zhang et al. 2021). Previous studies using *Arabidopsis thaliana* showed that the upregulation of the expression of senescence-associated genes (*SAG*s) during leaf senescence is controlled by tens of transcription factors (TFs), including the WRKY domain-containing transcription factors (WRKY TFs), as well as NAM, ATAF1/2, and CUC2 transcription factors (NAC TFs) (Weaver et al. 1998, Balazadeh et al. 2008, Kim et al. 2013). However, much remain to be known regarding the molecular mechanisms underlying the regulation of leaf senescence.

Actin microfilaments (AFs) are vital for many cellular functions such as intracellular trafficking, maintenance of cell shape, and cell division. The structures and dynamics of AFs are regulated by various actin-binding proteins (ABPs) (Borisy and Svitkina 2020, Paavilainen et al. 2004). In particular, ACTIN DEPOLYMERIZING FACTOR (ADF), one of ABPs, is involved in actin destabilization by depolymerizing and cleaving AFs (Ruzicka et al. 2007, Inada 2017). *A. thaliana* possesses 11 *ADF* genes that are classified into four subclasses based on similarities in their amino acid sequences (Ruzicka et al. 2007, Inada 2017). Subclass I *ADF*s, including *ADF1*, *-2*, *-3*, and *-4* are constitutively expressed in both vegetative and reproductive tissues but not in pollen. The level of expression of *ADF3* is the highest among members of subclass I *ADF*s. In turn, *ADF1* and *ADF2* show moderate expression, while *ADF4* shows only weak expression (Ruzicka et al. 2007).

In *A. thaliana*, subclass I ADFs are involved in the regulation of many physiological mechanisms, including plant growth and responses to both abiotic and biotic stressors. For example, *ADF1* knockdown transgenic lines exhibit elongated hypocotyls and roots (Dong et al. 2001). *ADF1* knockout mutants (*adf1*) exhibit enhanced plant growth under high-temperature conditions (Wang et al. 2023) and suppression of survival in response to salt stress (Wang et al. 2021). RNA interference (RNAi) line in which *ADF2* expression is suppressed shows inhibition of infection by the root-knot nematode *Meloidogyne incognita* (Clement et al. 2009). *ADF3* knockout mutant (*adf3*) are more susceptible to the nematode *Aphelenchoides besseyi* (Ding et al. 2022). *ADF4* knockout mutant (*adf4*) exhibits hypocotyl elongation in the dark and increases survival rates under osmotic conditions (Yao et al. 2022). *adf4* becomes more susceptible to bacterial pathogen *Pseudomonas syringae* DC3000 pv. tomato harvoring AvrPhpB (Tian et al. 2009, Porter et al. 2012). Previously, we have shown that both *adf4* and transgenic lines in which expressions of all members of subclass I *ADF*s are downregulated (*ADF1-4Ri*) show increased plant size and enhanced endoreplication (Inada et al. 2021). We also showed that *adf4* and *ADF1-4Ri* increase resistance against adapted-powdery mildew fungus *Golovinomyces orontii*, and that the nuclear localization of ADF4 is important for the response to *G. orontii* (Inada et al. 2016). Our recent report showed that *adf4* and *ADF1-4Ri* alter nuclear organization and expression of many genes (Matsumoto et al. 2023).

Here, we report a previously unknown function of subclass I ADFs in the regulation of leaf senescence. We found that *adf4* and *ADF1-4Ri* showed acceleration of both dark-induced and age-dependent leaf senescence. Further, qRT-PCR analysis revealed that the expression of *SAG*s was upregulated in *adf4* and *ADF1-4Ri* at earlier times in age-dependent leaf senescence than in wild-type Col-0 plants. Additionally, the expression of *ADF4* was downregulated during age-dependent leaf senescence. Complementation analysis indicated that the nuclear localization of ADF4 is important in regulating leaf senescence. This is the first study to demonstrate the involvement of subclass I ADFs in the regulation of leaf senescence.

## RESULTS

### Dark-induced leaf senescence was accelerated in *adf4*

In a previous study, we performed a microarray analysis using mature leaves of 4-week old wild-type Col-0, *adf4,* and *ADF1-4Ri* plants. This analysis showed that the expression of many genes was significantly altered in both *adf4* and *ADF1-4Ri* compared to Col-0 (Matsumoto et al. 2023). Upon close examination of the microarray data, we found that the expression of leaf senescence-associated genes was upregulated in *ADF1-4Ri* (Fig. S1), suggesting an acceleration of leaf senescence.

To assess this possibility, we first tested the progression of dark-induced senescence in Col-0, *adf3*, *adf4,* and *ADF1-4Ri* plants grown at 23°C under short-day conditions (8 h light; 16 h dark). The first and second leaves of 21-day-old plants were excised, placed in the incubation medium in the dark for 4 days, following a previously reported protocol (Zhang et al. 2018). Before dark treatment, the first and second leaves of 21-day-old *ADF1-4Ri* exhibited slight yellowing compared to those of Col-0, *adf3*, and *adf4* (Fig. 1A). There was no significant difference in chlorophyll content of those leaves of Col-0, *adf3,* and *adf4*, while *ADF1-4Ri* leaves exhibited lower chlorophyll content than Col-0 leaves (Fig. 1B). After 4 days of dark treatment, *adf4* and *ADF1-4Ri* showed a significant reduction in chlorophyll content compared to Col-0, whereas the chlorophyll content of dark-treated *adf3* was comparable to that of Col-0 (Fig. 1C, D). Based on these results, we concluded that dark-induced senescence was accelerated in *adf4*.

**Figure 1.**
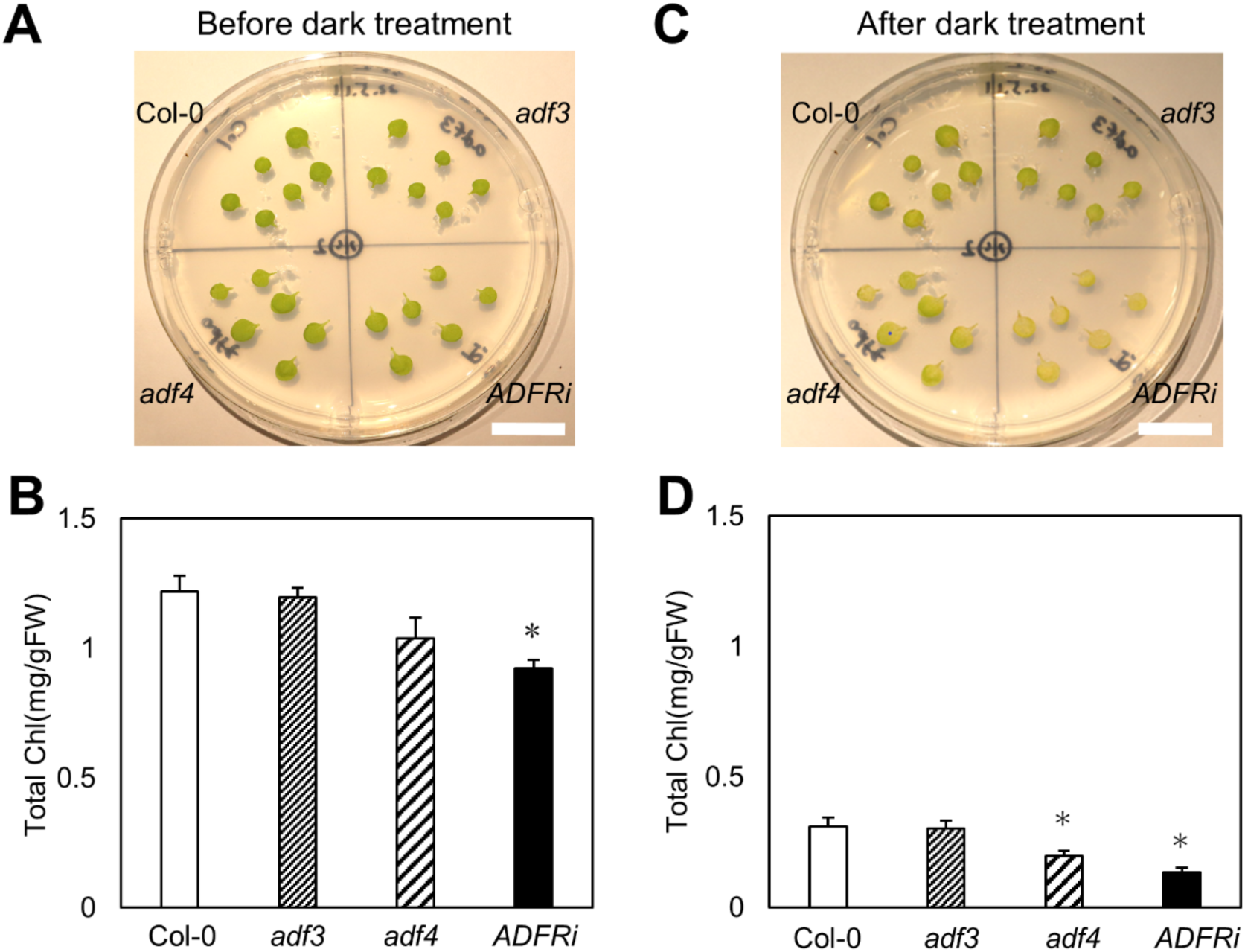
Loss of *ADF4* shows early leaf senescence when treated in the dark. (**A**) The first and the second leaves of 21-day-old Col-0, *adf3*, *adf4*, and *ADF1-4Ri* (*ADFRi*) grown at 23°C under short-day conditions (8 h light; 16 h dark). (**B**) Total chlorophyll content of the first and the second leaves of 21-day-old plants. (**C**) The first and the second leaves of Col-0, *adf3*, *adf4*, and *ADF1-4Ri* (*ADFRi*) treated in the dark for 4 days. (**D**) Total chlorophyll content of the first and the second leaves treated in the dark for 4 days. Bars in **A** and **C** indicate 1 cm. For **B** and **D**, 3 independent experiments were performed and the average values were shown. Error bars indicate standard deviation (SD). Comparison with Col-0 was performed by Student’s *t*-test. Asterisks indicate *P*<0.05.

Four independent lines were established for *ADF1-4Ri* (#1-4, #2-1, #3-2, #4-2; Tian et al. 2009), in which the expression of all members of subclass I *ADF* is suppressed by RNAi. We used *ADF1-4Ri*#2-1 for the experiments described above. To confirm the lower chlorophyll content in *ADF1-4Ri*#2-1 leaves, we examined three other lines of *ADF1-4Ri*, *ADF1-4Ri* #1-4, #3-2, and #4-2. The first and second leaves of 21-day-old *ADF1-4Ri* #1-4 and #3-2, but not #4-2, had lower chlorophyll content than the leaves of Col-0 plants (Fig. S2A, B). However, after 4 days of dark treatment, all three *ADF1-4Ri* lines showed significantly lower chlorophyll content than Col-0 plants (Fig. S2C, D). Therefore, we used *ADF1-4Ri*#2-1 in subsequent experiments as a representative *ADF1-4Ri* line.

### Age-dependent leaf senescence was accelerated in *adf4* and *ADF1-4Ri*

The fact that the chlorophyll content in the first and second leaves of 21-day-old *ADF1-4Ri* was lower than that in Col-0 plants, even before dark treatment (Fig. 1A, B), might indicate either reduced chlorophyll accumulation during leaf development or accelerated chlorophyll degradation due to accelerated leaf senescence in *ADF1-4Ri*. To investigate these possibilities, we determined the chlorophyll content of the first and second leaves at the early and late leaf developmental stages. At age 14 days, plants grown at 23°C under short-day conditions had started to develop the third and fourth leaves, while the first and second leaves were expanding (Fig. 2A). The chlorophyll content in the first and second leaves of 14-day-old *ADF1-4Ri* plants was comparable to that of Col-0 plants, indicating no impairment in chlorophyll accumulation during leaf development (Fig. 2B). On the other hand, 35-day-old plants showed the first and second leaves had started to show slight yellowing in *ADF1-4Ri* (Fig. 2C). Chlorophyll contents in the first and second leaves of 35-day-old *ADF1-4Ri* were significantly lower than those in Col-0 plants (Fig. 2D). Similarly, we tested other lines of *ADF1-4Ri* and found that *ADF1-4Ri* #1-4 and #3-2 lines, but not *ADF1-4Ri* #4-2, also showed reduced chlorophyll contents in the first and second leaves of 35-day-old plants, strongly suggesting that age-dependent leaf senescence was accelerated in *ADF1-4Ri* plants (Fig. S3A, B).

**Figure 2.**
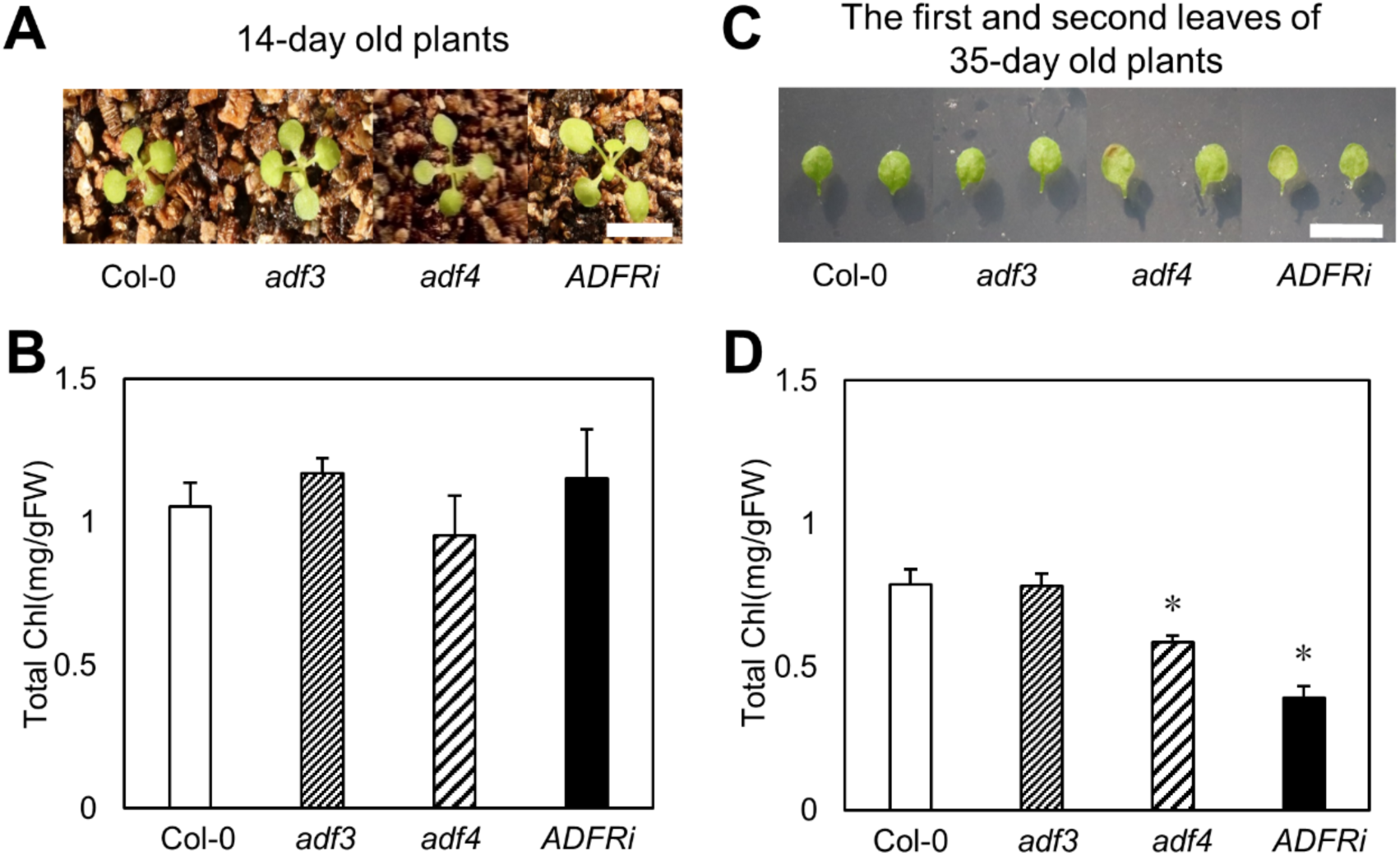
*adf4* and *ADF1-4Ri* show early age-dependent leaf senescence. (**A**) 14-day-old Col-0, *adf3*, *adf4*, and *ADF1-4Ri* (*ADFRi*) grown at 23°C under short-day conditions. (**B**) Total chlorophyll content of the first and the second leaves of 14-day-old plants. (**C**) The first and second leaves excised from 35-day-old Col-0, *adf3*, *adf4*, and *ADF1-4Ri* (*ADFRi*) grown at 23°C under short-day conditions. (**D**) Total chlorophyll content of the first and the second leaves of 35-day-old plants. Bars in **A** and **C** indicate 1 cm. For **B** and **D**, 3 independent experiments were performed and the average values were shown. Error bars indicate SD. Comparison with Col-0 was performed by Student’s *t*-test. Asterisks indicate *P*<0.05.

Additionally, we analyzed the chlorophyll contents of *adf3* and *adf4* plants during age-dependent senescence. The first and second leaves of 14-day-old seedlings of both mutants showed no significant differences in chlorophyll content compared to Col-0 plants (Fig. 2A, B). The chlorophyll content in the first and second leaves of 35-day-old *adf3* mutant plants was also comparable to that of Col-0 plants. However, the leaves of 35-day-old *adf4* mutant showed lower chlorophyll contents than those of Col-0 (Fig. 2C, D). Another *adf4* T-DNA insertion line (*adf4-2*) also showed lower chlorophyll content than Col-0 when the first and second leaves of 35-day-old plants were analyzed (Fig. S3C, D), indicating that ADF4 plays a major role in the regulation of age-dependent leaf senescence.

In addition to programmed leaf senescence, several stresses on photosynthetic machinery induces chlorophyll degradation (Hörtensteiner and Kräutler 2011). To assess whether the accelerated chlorophyll degradation in *adf4* and *ADF1-4Ri* is associated with disruption of photosynthetic machinery that triggers leaf senescence, we examined the maximal quantum efficiency of photosystem II (*F*_V_/*F*_M_), which reflects the functionality of photosystem II and thus the photosynthetic capability of leaves, with a chlorophyll fluorescence imager. For this analysis, we used true leaves from 35-day-old plants grown under the short-day condition (Fig. 3A, B). In all plants tested, *F*_V_/*F*_M_ values of the 13th and 14th leaves reached to around 0.8, suggesting that chloroplasts were fully functional in these young leaves (Fig. 3C). Similar results were observed in other leaves (Fig. 3B). Even in the first and second leaves, *F*_V_/*F*_M_ levels were not significantly decreased as compared to those in the 13th and 14th leaves in Col-0, *adf3*, and *adf4*. As an exception, the first and second leaves of *ADF1-4Ri* showed a slight decrease in *F*_V_/*F*_M_, but the level around 0.77 was sufficiently high to maintain the photosynthetic electron transport in chloroplasts. The data indicate that *adf4* and *ADF1-4Ri* plants accelerate chlorophyll degradation in developed leaves while maintaining the photosynthetic functionality. Nevertheless, we found that 11.1% and 16.7% of the first and second leaves of the *adf4* and *ADF1-4Ri* plants, respectively, appeared withered and had almost no chlorophylls, whereas Col-0 and *adf3* had no such withered true leaves at the 35-day-old growth stage (Fig. 3D), confirming the accelerated leaf senescence in *adf4* and *ADF1-4Ri*. Taken together, our data suggest that the intrinsic senescence program, but not the disruption of photosynthetic machinery, is accelerated in *adf4* and *ADF1-4Ri* plants.

**Figure 3.**
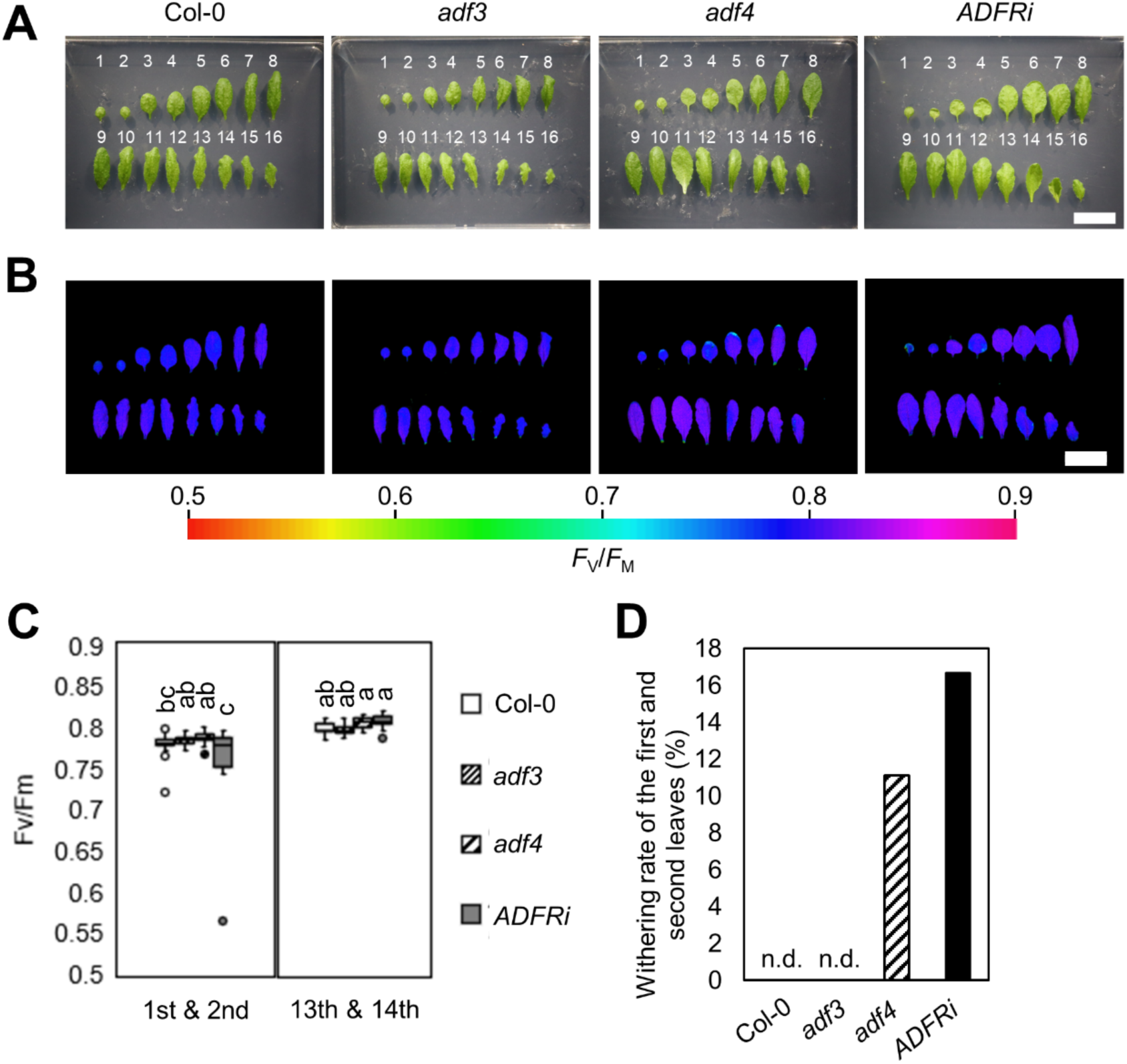
*adf4* and *ADF1-4Ri* maintain photosystem II functionality during leaf senescence. (**A**) Ture leaves detached from 35-day-old Col-0, *adf3*, *adf4*, and *ADF1-4Ri* (*ADFRi*) grown at 23°C under short-day conditions. Leaf numbers are shown in the image for each line. (**B**) Images of maximum quantum efficiency of photosystem II (*F*_V_/*F*_M_) in leaves at different developmental stages. Leaves are the same as shown in (**A**). Bars in **A** and **B** indicate 2 cm. Representative images from 9 independent experiments were shown. The color represents the value of *F*_V_/*F*_M_ in the color scale from 0.5 to 0.9. (**C**) *F*_V_/*F*_M_ levels in the first and second leaves (1st & 2nd) and those in the 13th and 14th leaves (13th & 14th). Each box indicates the interquartile range with the midline representing the median value. The whiskers extend to the lowest/highest values that is no >1.5 times the interquartile range, with outliers shown by dots (n = 18 for Col-0 and *adf3*, n = 16 for *adf4*, and n = 15 for *ADF1-4Ri*). The lowercase letters indicate statistically significant differences between each line (*P* < 0.05, 1-way ANOVA and Tukey’s post hoc honestly significant difference test). (**D**) Withering rate of the first and second leaves (n = 18 for each line). Leaves that had almost no chlorophyll fluorescence in the *F*_V_/*F*_M_ analysis were defined as withered one. n.d., not detected.

### *act2/8* and *act7* mutants did not show accelerated leaf senescence

*A. thaliana* genome contains eight *ACT* genes (McDowell et al. 1996a, Meagher et al. 1999). Among these, *ACT2*, *-7*, and *-8* are expressed in vegetative organs (An et al. 1996, McDowell et al. 1996b). In particular, ACT2 and ACT8 differ by only one amino acid and exhibit functional redundancy (Kandasamy et al. 2009, Kandasamy et al. 2010). Previous studies showed that *ACT* mutants exhibit phenotypes similar to those of *adf4* and *ADF1-4Ri*; the double mutant for *ACT2* and *ACT8* (*act2/8*) shows increased leaf area and enhanced endoreplication (Inada et al. 2021); chromocenter size is altered in *adf4* and *ADF1-4Ri*, as well as in *act7* (Matsumoto et al. 2023). Therefore, we examined whether the progression of leaf senescence was also altered in *ACT* mutants.

To examine the progression of dark-induced senescence in *ACT* mutants, the first and second leaves of 21-day-old Col-0, *act2/8,* and *act7* plants were treated in the dark, and the leaf chlorophyll contents were measured. The results showed that, before (Fig. 4A, B) and after (Fig. 4C, D) dark treatment, the chlorophyll contents of *act2/8* and *act7* were comparable to those of Col-0 plants. To investigate whether the loss of *ACT* accelerated age-dependent leaf senescence, the chlorophyll contents of the first and second leaves of 35-day-old Col-0, *act2/8,* and *act7* plants were measured. We found no significant difference in chlorophyll contents among these phenotypes (Fig. 4E, F). Thus, neither dark-induced nor age-dependent senescence was accelerated in *act2/8* or *act7* mutant plants.

**Figure 4.**
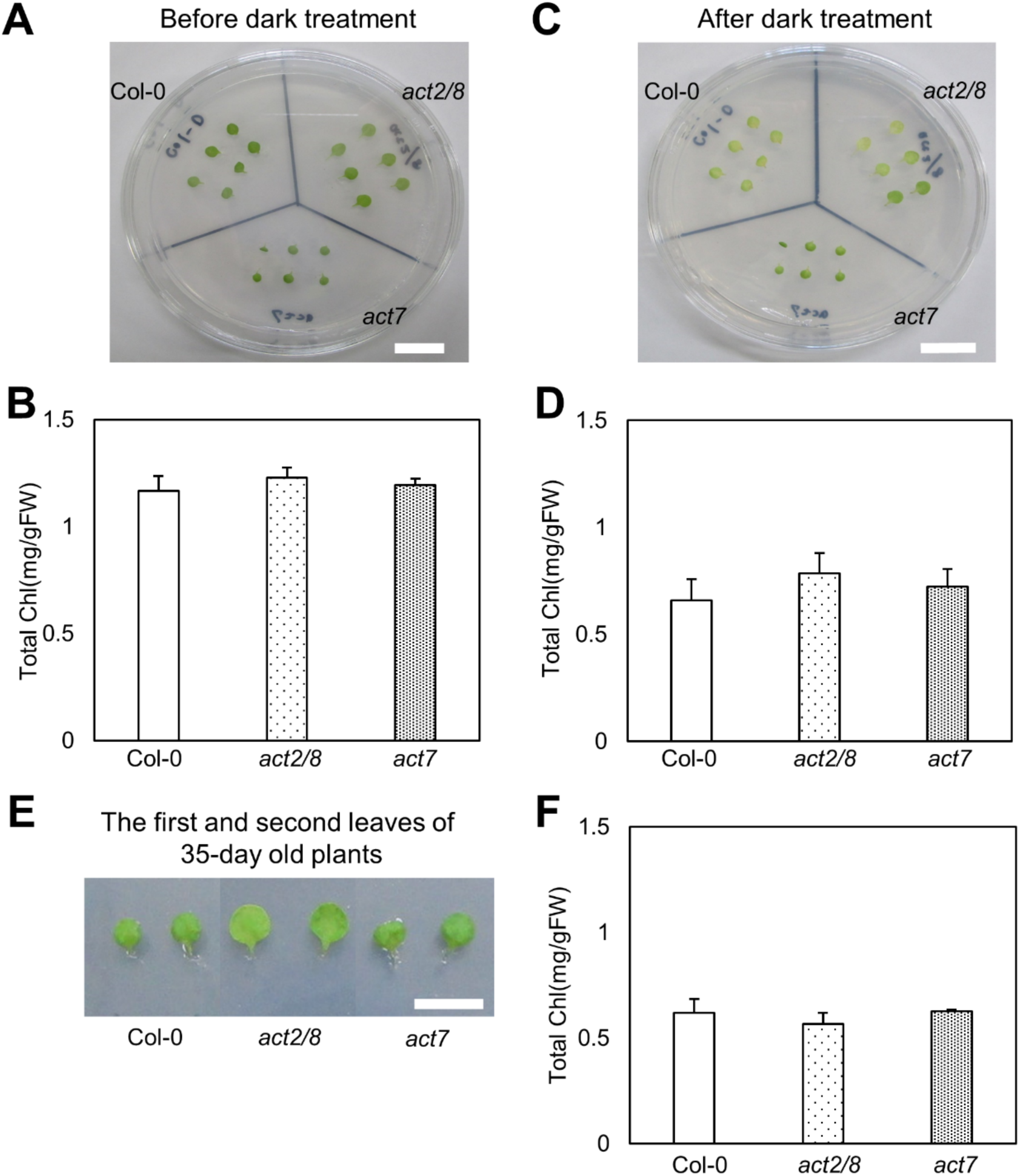
Neither dark-induced nor age-dependent leaf senescence was affected in *act2/8* and *act7.* (**A**) The first and the second leaves of 21-day-old Col-0, *act2/8*, and *act7* grown at 23°C under short-day conditions. (**B**) Total chlorophyll content of the first and the second leaves of 21-day-old plants. (**C**) The first and the second leaves of Col-0, *act2/8*, and *act7* treated in the dark for 4 days. (**D**) Total chlorophyll content of the first and the second leaves treated in the dark for 4 days. (**E**) The first and the second leaves of 35-day-old Col-0, *act2/8*, and *act7* grown at 23°C under short-day conditions. (**F**) Total chlorophyll content of the first and the second leaves of 35-day-old plants. Bars in **A**, **C,** and **E** indicate 1 cm. For **B**, **D**, and **F**, 3 independent experiments were performed, and the average values were shown. Error bars indicate SD. Comparison with Col-0 was performed by Student’s *t*-test. Asterisks indicate *P*<0.05.

### The expression of *SAG*s was upregulated in *adf4* and *ADF1-4Ri*

The expression of several *SAG*s was upregulated during dark-induced and age-dependent leaf senescence (Weaver et al. 1998, Kim et al. 2013, Zhang et al. 2018). To understand the molecular basis of accelerated leaf senescence in *adf4* and *ADF1-4Ri*, we analyzed the expression of *SAG*s during age-dependent leaf senescence by qRT-PCR.

Four *SAG*s (At2g29350, At4g22920, At5g13800, and At4g23810) were selected for qRT-PCR analysis. These genes were identified as genes upregulated during age-dependent leaf senescence by transcriptome profiling during leaf senescence (Guo et al. 2004). At2g29350 encodes the SENESCENCE-ASSOCIATED GENE 13 (SAG13) protein, a widely conserved protein with unknown function (Lohman et al. 1994, Dhar et al. 2020). *A. thaliana SAG13* knockout mutants show delayed dark-induced leaf senescence (Dhar et al. 2020). At4g22920 encodes an Mg-dechelatase called STAY GREEN1 (SGR1) or NON-YELLOWING1 (NYE1), which catalyzes the initial step of the chlorophyll a degradation pathway during leaf senescence, namely the release of Mg^2+^ from chlorophyll *a* to form pheophytin *a* (Shimoda et al. 2016). In *A. thaliana*, overexpression of *SGR1/NYE1* enhanced chlorophyll degradation, whereas the loss of function of this gene impaired chlorophyll degradation during dark-induced leaf senescence (Ren et al. 2007, Sakuraba et al. 2012). At5g13800 encodes PHEOPHYTINASE (PPH), which removes the phytol chain from pheophytin *a*, resulting in pheophoribide *a*, an intermediate product of the chlorophyll degradation pathway. *PPH* knockout mutants exhibit delayed dark-induced leaf senescence (Schelbert et al. 2009). At4g23810 encodes the WRKY domain-containing TF 53 (WRKY53). WRKY is a TF family with a zinc-finger structure that regulates stress responses, plant development, and leaf senescence (Miao et al. 2004). *A. thaliana WRKY53* RNAi lines showed delayed age-dependent leaf senescence (Miao et al. 2004).

Col-0, *adf3, adf4*, and *ADF1-4Ri* plants were grown at 23°C under short-day conditions, and the first and second leaves of 21-, 28-, and 35-day-old plants were used for RNA extraction. The expression of all tested *SAG*s in Col-0 plants was upregulated as age-dependent senescence progressed (Fig. S4A-D). Although we did not observe an early yellowing phenotype in *adf3* (Fig. 2C, D), we found that the expression of *SAG*s, *SAG13, SGR1*, and *PPH*, but not *WRKY53*, was upregulated in *adf3* in 21-day-old (Fig. 5A, D, G) and 28-day-old (Fig. 5B, E, H) plants compared with Col-0. Moreover, the expression of *WRKY53* at 35-day-old plants also was upregulated in *adf3* compared with Col-0 (Fig. 5L). In *adf4*, the expression of *SGR1*, *PPH*, and *WRKY53*, but not *SAG13*, was upregulated at age 21 days (Fig. 5D, G, J). At age 28 days, *PPH* showed higher expression in *adf4* compared to Col-0 (Fig. 5H). The expression of *SAG13* and *SGR1* in *adf4* was then downregulated compared to Col-0 at age 35 days (Fig. 5C, F). Similarly, *ADF1-4Ri* showed upregulation of many *SAG*s, compared to Col-0; in particular, the expression of *SAG13, SGR1*, and *PPH* was upregulated at age 21 and 28 days (Fig. 5A, B, D, E, G, H). Moreover, the expression of *SAG13* at age 35 days (Fig. 5C) and that of *WRKY53* at age 28 days (Fig. 5K) also increased in *ADF1-4Ri*, compared to Col-0 plants at each age.

**Figure 5.**
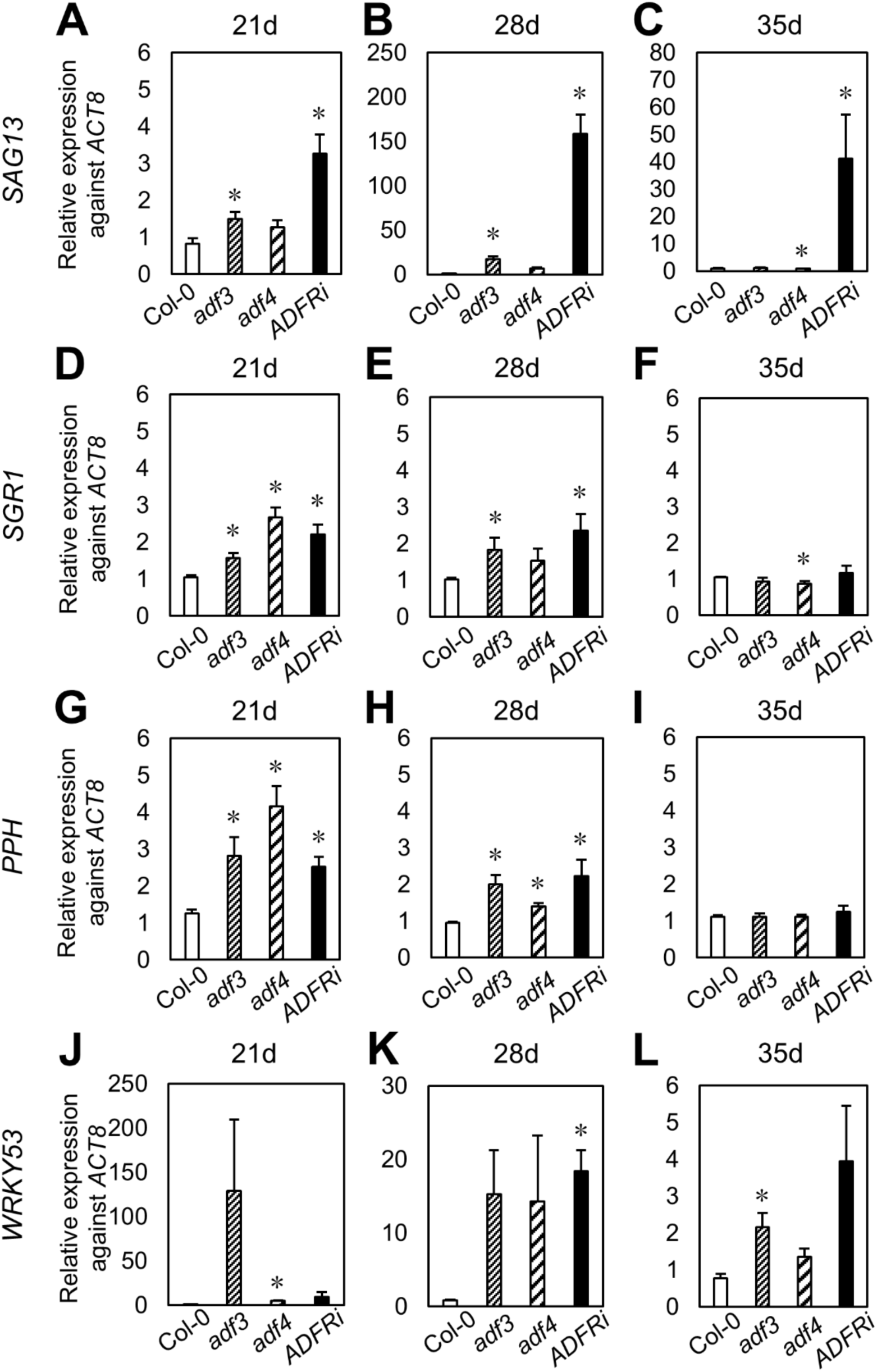
The expression of *SAG*s was upregulated in *adf4* and *ADF1-4Ri*. The expression of *SAG13* (**A**-**C**), *SGR1* (**D-F**), *PPH* (**G-I**), and *WRKY53* (**J-L**) in the first and the second leaves of Col-0, *adf3*, *adf4* and *ADF1-4Ri* at 21- (**A, D, G, J**), 28- (**B, E, H, K**), and 35- (**C, F, I, L**) days. For normalization, *ACT8* (At1g49290) was used as the internal control. The average value was obtained from more than two biological replicates, one biological replicate containing three technical replicates. Error bars indicate SD. Comparison with Col-0 was performed by Student’s *t*-test. Asterisks indicate *P*<0.05.

Based on these results, we concluded that the onset of leaf senescence occurred earlier in *adf4* and *ADF1-4Ri* than in Col-0 plants.

### The expression of *ADF4* was downregulated during age-dependent leaf senescence

To investigate whether the expression of subclass I *ADF*s is altered during age-dependent leaf senescence, qRT-PCR was performed using RNA extracted from the first and second leaves of 14-, 21-, 28- and 35-day-old Col-0 plants. The first and second leaves of Col-0 were fully expanded at age 14 days, and no yellowing was visible at 21, 28, or 35 days. *SAG13* was used as a leaf senescence marker.

The expression of *SAG13* was too low to be quantified on day 14 but it was quantified on day 21, and increased on days 28 and 35 (Fig. 6A). The expression of both *ADF1* and *ADF2* decreased on day 21, increased on day 28, and then decreased again on day 35 (Fig. 6B, C). The expression of *ADF3* also showed a transient increase after 28 days (Fig. 6D). In contrast, the expression of *ADF4* was downregulated as age-dependent senescence progressed (Fig. 6E). These results suggest that among subclass I ADFs, ADF4 plays a major role in the regulation of leaf senescence.

**Figure 6.**
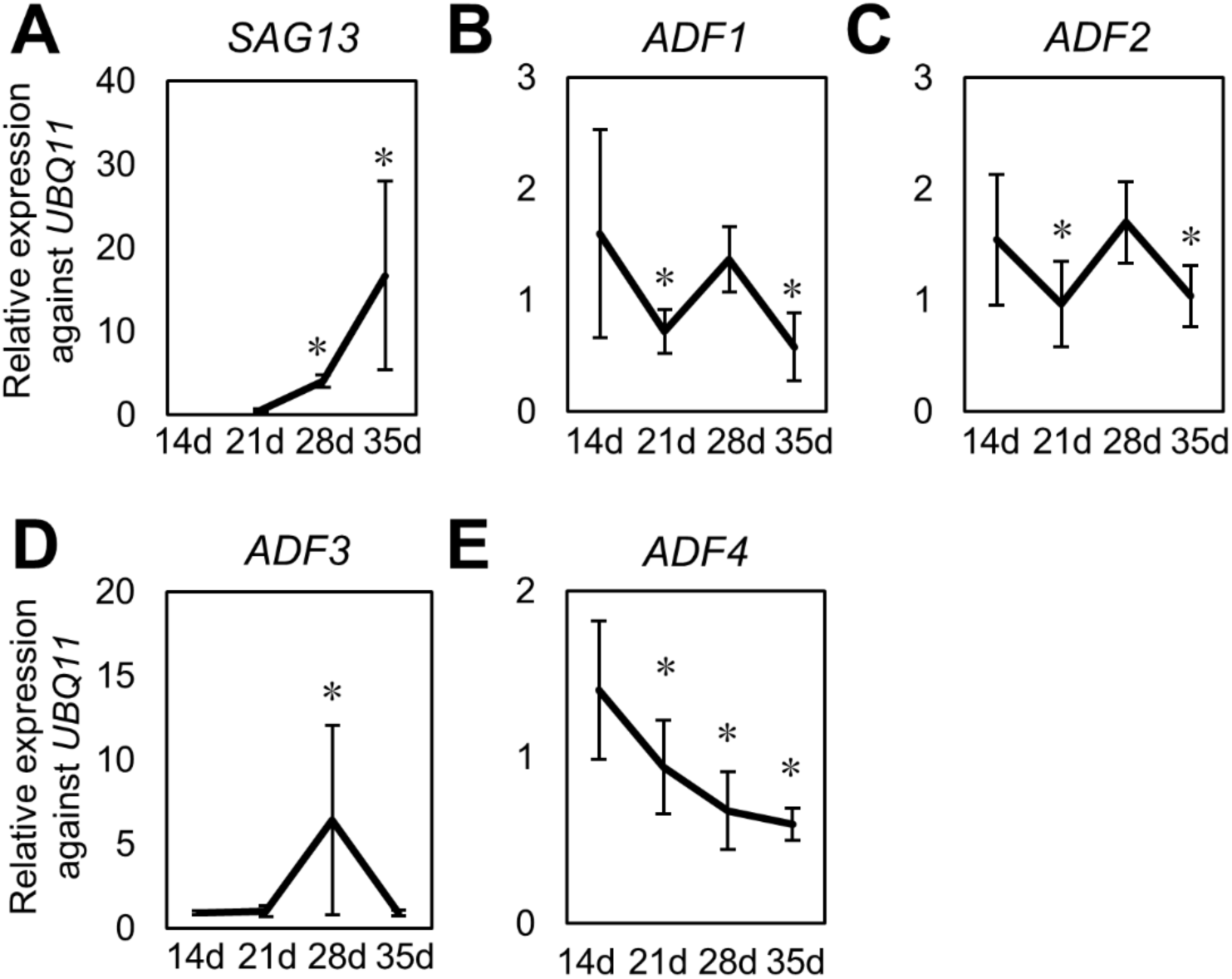
The expression of *ADF4* was downregulated in Col-0 during age-dependent leaf senescence. The expression of *SAG13* (**A**), *ADF1* (**B**), *ADF2* (**C**), ADF3 (**D**), and *ADF4* (**E**) in the first and the second leaves of plants at 14-, 21-, 28- and 35-days. For normalization, *UBQ11* (At4g050050) was used as the internal control. The average values were obtained from more than two biological replicates, one biological replicate containing three technical replicates. Error bars indicate SD. Comparison with expression at 21-days (**A**) and at 14-days (**B-E**) was performed by Student’s *t*-test. Asterisks indicate *P*<0.05.

### Nuclear localization of ADF4 was important for the regulation of age-dependent leaf senescence

ADF4 co-localizes with AFs in the cytoplasm and localizes to the nucleus as well (Ruzicka et al. 2007, Tian et al. 2009, Porter et al. 2012, Inada et al. 2016). Previous studies showed that nuclear localization of ADF4 is important for the response to the powdery mildew fungus *G. orontii* (Inada et al. 2016). The *adf4* phenotype showed increased resistance to *G. orontii* and was complemented by the expression of ADF4 fused to GFP (ADF4-GFP), which localized to the nucleus in addition to the cytoplasmic AFs. In contrast, the expression of ADF4-GFP fused with a nuclear exporting signal (NES), which co-localized with cytoplasmic AFs but did not localize to the nucleus, did not complement the increased *adf4* resistance to *G. orontii* (Inada et al. 2016).

To investigate whether the nuclear localization of ADF4 is important for the regulation of age-dependent leaf senescence, we measured the chlorophyll content in the first and second leaves of Col-0, *adf4,* ADF4-GFP/*adf4,* and ADF4-GFP-NES/*adf4* at age 35 days, when the leaves of *adf4* started to show a reduction in chlorophyll content compared to Col-0 (Fig.2C, D). Meanwhile, the chlorophyll content of ADF4-GFP/*adf4* was comparable to that of Col-0 plants (Fig. 7A, B), indicating that the early leaf senescence phenotype of *adf4* was complemented by the expression of ADF4-GFP. In contrast, the chlorophyll content of ADF4-GFP-NES/*adf4* was lower than that of the Col-0. These results indicated that nuclear localization of ADF4 is important for the regulation of leaf senescence.

**Figure 7.**
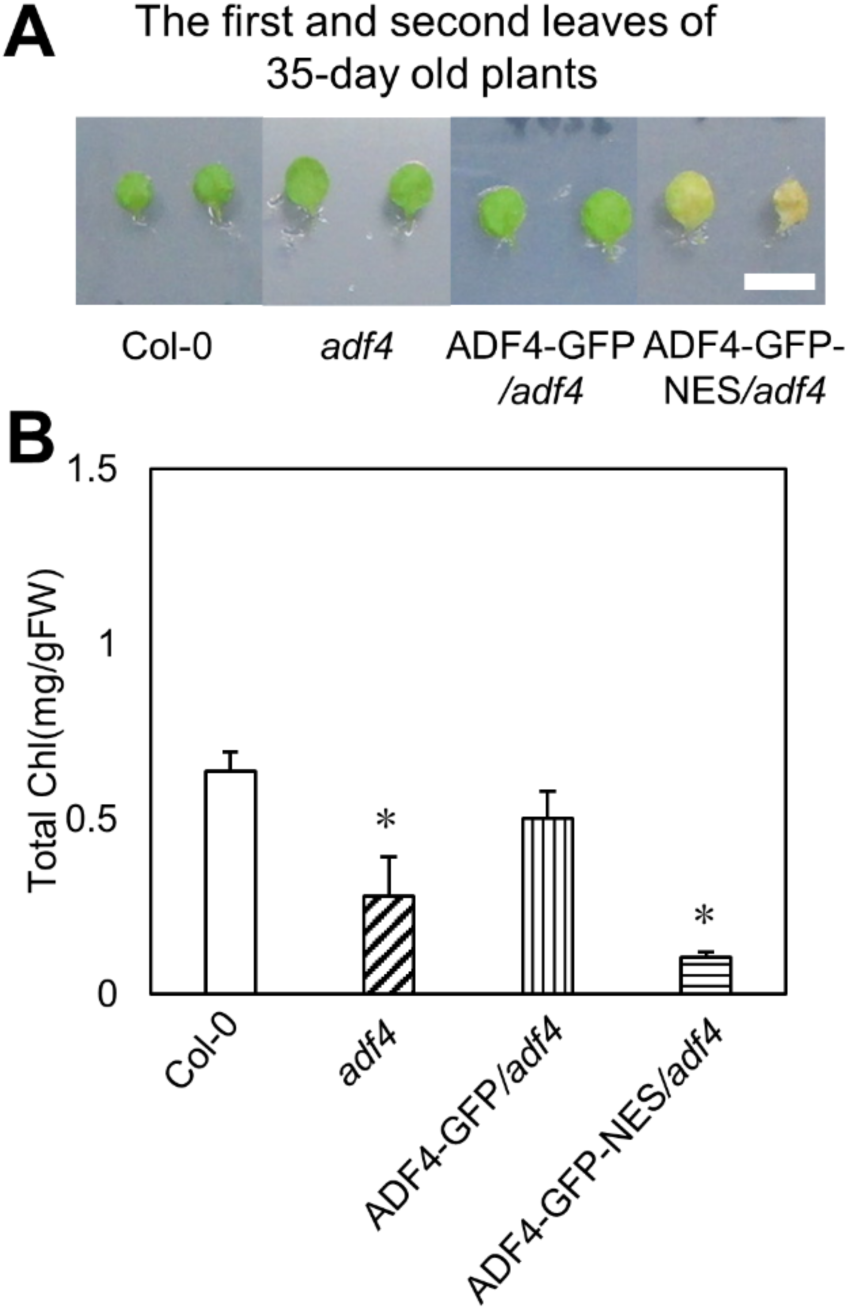
Nuclear localization is important for ADF4 function in regulation of age- dependent leaf senescence. (**A**) The first and the second leaves of 35-day-old Col-0, *adf4*, ADF4-GFP/*adf4*, and ADF4-GFP-NES/*adf4* grown at 23°C under short-day conditions. (**B**) Total chlorophyll content of the first and the second leaves of 35-day-old plants. Bar in **A** indicates 1 cm. For **B**, 3 independent experiments were performed, and the average values were shown. Error bars indicate SD. Comparison with Col-0 was performed by Student’s *t*-test. Asterisks indicate *P*<0.05.

## DISCUSSION

In this study, we found that *A. thaliana* subclass I ADFs are previously unknown regulators of leaf senescence. Both dark-induced and age-dependent leaf senescence were accelerated in *adf4* and *ADF1-4Ri* compared with Col-0. Chlorophyll fluorescence analysis suggests that not chloroplast dysfunction, but induction of intrinsic mechanisms would accelerate leaf senescence in *adf4* and *ADF1-4Ri*. Further, *ADF1-4Ri* showed an earlier onset of leaf senescence compared to *adf4* mutant; in *ADF1-4Ri*, chlorophyll content of the first and second leaves was already lower than that in Col-0 at age 21 days, while the chlorophyll content in *adf4* at this age was comparable to that in Col-0. In contrast, *act2/8* and *act7* did not show accelerated leaf senescence. Additionally, we found that the expression of *SAG*s was upregulated in *adf4* and *ADF1-4Ri*, compared with Col-0. Although *adf3* did not show accelerated degradation of chlorophyll during dark-induced or age-dependent leaf senescence, the expression of *SAG*s was upregulated earlier than that in Col-0, suggesting *ADF3* is also involved in the suppression of *SAG* expression. In addition, the expression of *ADF4*, but not that of *ADF1*, *ADF2*, or *ADF3*, was downregulated during age-dependent leaf senescence. Complementation analysis using ADF4-GFP/*adf4* and ADF4-GFP-NES/*adf4* indicated the importance of the nuclear localization of ADF4 for the regulation of leaf senescence. Based on these results, we conclude that subclass I ADFs, especially ADF4, are previously unknown regulators of leaf senescence and that nuclear localization of ADF4 is important for the suppression of leaf senescence.

*ADF1-4Ri* showed a much earlier onset of leaf senescence than *adf4*, as indicated by the fact that the chlorophyll content at age 21 days had already decreased in *ADF1-4Ri* (Fig.1A, B), suggesting that subclass I ADFs function redundantly in the regulation of leaf senescence. Among the four members of subclass I *ADF*s, *ADF3* showed the greatest expression level in many tissues, whereas the expression of *ADF4* showed the lowest (Ruzicka et al. 2007). In contrast to expression levels, the acceleration of leaf senescence was more striking in *adf4* than in *adf3*. Among the four subclass I *ADF*s, only *ADF4* showed a strong correlation between changes in expression and leaf senescence. These results indicate that ADF4 is the main regulator of leaf senescence among subclass I ADFs, although our observation that expression of several *SAG*s was upregulated in *adf3* at ages 21 and 28 days suggests that ADF3 also functions in the regulation of leaf senescence. Of note, the expression of *SGR1* and *PPH*, whose translation products catalyze the essential steps of the chlorophyll degradation pathway (Schelbert et al. 2009., Shimoda et al. 2016), was upregulated in *adf4* and *ADF1-4Ri*. Thus, increased expression of these genes may contribute to the earlier loss of chlorophyll in both plants. However, *adf3*, which showed no accelerated chlorophyll degradation, also had increased expression of *SGR1* and *PPH*. The data suggest that the accelerated chlorophyll degradation in *adf4* and *ADF1-4Ri* would include complex mechanisms other than the increased *SGR1* and *PPH* expression.

The suppression of *ADF* expression leads to the stabilization of AFs, which causes a decrease in the amount of monomeric actin and alters AF organization (Dong et al. 2001). To investigate whether a decrease in the amount of monomeric actin due to the loss of *ADF* is the primary cause of accelerated leaf senescence, we examined whether mutations in *ACT* genes, which reduce total actin levels (Kandasamy et al. 2009), concomitantly accelerate leaf senescence. The results showed that leaf senescence progression in *ACT* mutants was comparable to that in Col-0 plants (Fig. 4), suggesting that the decrease in the amount of monomeric actin in *adf4* and *ADF1-4Ri* was not associated with the regulation of leaf senescence. In contrast, alterations in the AF organization are reportedly involved in the regulation of leaf senescence. Interestingly, myosins are actin-based molecular motors, and the inactivation of *A. thaliana* class XI myosins is known to affect AF organization and the progression of leaf senescence (Ojangu et al. 2018). In this study, triple knockout (3KO) mutants of class XI myosin, *XI1*, *XI2*, and *XIK*, showed a loss of chlorophyll content and strong upregulation of *SAG13* during dark-induced senescence (Ojangu et al. 2018). *xi1 xi2 xik* 3KO plants showed thinner AF arrays than Col-0 plants. While AFs in the Col-0 midvein epidermis showed longitudinal organization, *xi1 xi2 xik* 3 KO showed transverse AFs before age-dependent senescence (Peremyslov et al. 2010, Ojangu et al. 2018). Furthermore, treatment with the AF- destabilizing drug latrunculin B (LatB) upregulated the expression of *SAG13* (Ojangu et al. 2018). Thus, alteration in AF organization due to ADF loss may be involved in the regulation of leaf senescence.

Our results showed that ADF4-GFP, but not ADF4-GFP-NES, complemented the early senescence phenotype of *adf4*, indicating that nuclear localization of ADF4 is important for the regulation of leaf senescence. Using the same lines, we previously showed that the nuclear localization of ADF4 plays an important role in the response against powdery mildew fungus (Inada et al. 2016). Furthermore, we recently reported that subclass I ADFs regulate chromocenter organization and gene expression (Matsumoto et al. 2023). These results suggest that subclass I ADFs regulate leaf senescence by modulating the expression of *SAG*s in the nucleus. Nuclear AFs are reportedly involved in the regulation of chromatin organization and gene expression in mammalian cells (Fernandez et al. 2024). Various ABPs are located in the nucleus and are involved in the regulation of chromatin organization and gene expression through the regulation of nuclear AFs (Dingova et al. 2009, Kristó et al. 2016). In NIH3T3 cells, cultured cells derived from mice NIH/Swiss embryos, nuclear AFs are transiently formed when daughter cell nuclei are formed during mitotic cell division. Cofilin-1, a mammalian ADF, is involved in mitotic chromatin decomposition in daughter cells by promoting the destabilization of nuclear AFs (Baarlink et al. 2017). Formins, AF assembly activators, regulate the transient formation of nuclear AFs in response to serum treatment in NIH3T3 cells. Transiently formed AFs promote megakaryocytic acute leukemia (MAL) protein-serum response factor (SRF)- dependent transcription, thus promoting transcriptional activation in response to serum (Baarlink et al. 2013). The pro-oncogenic transcriptional co-activators YAP/TAZ promote tumorigenesis when cells are at high mechanics. YAP/TAZ activity is inhibited by its interaction with the switch/sucrose non-fermentable (SWI/SNF) chromatin remodeling complex. Nuclear AFs reportedly accumulate at high mechanics and bind to the SWI/SNF complex, consequently preventing the interaction between the SWI/SNF complex and YAP/TAZ, thus inducing YAP/TAZ transcriptional activity in human mammary epithelial cells (Chang et al. 2018). In plants, the formation of nuclear AFs has not been reported, whereas the presence of actin in the nucleus has been observed in *Allium cepa* (Cruz and Moreno 2009) and *A. thaliana* (Kandasamy et al. 2010). Our results that ADF functions in the nucleus, together with reports on nuclear AFs in cultured mammalian cells, suggest that *A. thaliana* subclass I ADFs are involved in the regulation of chromocenter organization and gene expression through the modulation of AFs in the nucleus.

Altogether, our results suggest that subclass I ADFs, particularly ADF4, are involved in the regulation of leaf senescence. Further investigation of ADF function in the regulation of leaf senescence will contribute to a thorough understanding of the mechanisms of control over leaf senescence and elucidate novel roles of ABPs in plant development.

## MATERIALS AND METHODS

### Plant materials and growth conditions

T-DNA insertional mutants *adf3* (Salk_139265) and *adf4-2* (Salk_ 121647) were obtained from Arabidopsis Biological Resource Center (ABRC). *adf4* (Garlic_823_A11.b.1b.Lb3Fa) and *ADF1-4Ri* were provided by Dr. Brad Day at Michigan State University (Tian et al. 2009). *act2/8*(*act2-1/act8-2*) and *act7* (SALK_13610) were provided by Dr. Tomohiro Takatsuka at Kanazawa University (Takatsuka et al. 2018). ADF4-GFP/*adf4* and ADF4-GFP-NES/*adf4* were produced previously (Inada et al. 2016).

Seeds were suspended in 0.1% agarose and vernalized at 4 °C for at least 1 day. Seeds after vernalization were sown in soil mixed at 3:1 vermiculite: Metromix and kept at high humidity for 1 week. Plants were grown in a growth chamber (LH-240S, NK Systems, Osaka, Japan) at 23°C under 8 h light:16 h dark photoperiod.

### Dark treatment

The first and second leaves were harvested from 3 individual plants at 21-day and placed on 5 mM MES solid medium (0.8% agarose, pH=5.7 with KOH). The MES solid medium covered with aluminum was kept in a growth chamber (LH-30CCFL-8CT, NK Systems, Osaka, Japan) at 23°C under 24 h dark for 4 days.

### Chlorophyll content measurement

The fresh weight of the first and second leaves was measured. Chlorophyll was extracted by treatment of leaves with 80% acetone. The absorbance of extracted chlorophyll was measured at 720, 646.8, and 663.2 nm (V-730BIO, JASCO, Tokyo, Japan). The sum of chlorophyll *a* and *b* content was determined by Lichtenthaler formula (Lichtenthaler 1987).

### *F*_V_/*F*_M_ determination

To determine *F*_V_/*F*_M_, leaves detached from 35-day-old plants were placed on the MES solid medium and dark-adapted for 20 min before measurement. By the use of an IMAGING-PAM MAXI fluorometer and ImagingWin software (Heinz Walz), the minimal (*F*_O_) and maximal chlorophyll fluorescence (*F*_M_) from leaves were determined under measuring light before and after saturating flash, respectively. *F*_V_/*F*_M_, values were calculated by the following equation: *F*_V_/*F*_M_ = (*F*_M_ − *F*_O_)/*F*_M_.

### qRT-PCR

The first and second leaves harvested from 4 individual plants were frozen in liquid N_2_. Total RNA was extracted using Sepasol®-RNAISugerG reagent, following the manufacturer’s protocol (NAKALAI TESQUE, Kyoto, Japan). The reverse transcription was carried out using the ReverTra Ace qPCR RT Master Mix with gDNA Remover (TOYOBO, Osaka, Japan) using 100 ng of total RNA according to the manual. qRT-PCR was carried out using THUNDERBIRD SYBR qPCR Mix (TOYOBO, Osaka, Japan) on 7300 Real-Time PCR System (Applied Biosystems, Massachusetts, USA). For normalization, *ACT8* (At1g49290) and *UBQ11* (At4g050050) were used as the internal control. The average value shown in the figures was obtained from more than two biological replicates, one biological replicate containing three technical replicates. Primer information is shown in Table S1.

## REFERENCES

An, Y-Q., McDowell, J.M., Huang, S., McKinney, E.C., Chambliss, S., and Meagher, R.B. (1996) Strong, constitutive expression of the *Arabidopsis ACT2/ACT8* actin subclass in vegetative tissues. Plant J. 10: 107–121.

Baarlink, C., Wang, H., and Grosse, R. (2013) Nuclear actin network assembly by formins regulates the SRF coactivator MAL. Science 340: 864–867.

Baarlink, C., Plessner, M., Sherrard, A., Morita, K., Misu, S., Virant, D. et al. (2017) A transient pool of nuclear F-actin at mitotic exit controls chromatin organization. Nat. Cell Biol. 19: 1389–1399.

Balazadeh, S., Riano-Pachon, D.M., and Mueller, R.B. (2008) Transcription factors regulating leaf senescence in *Arabidopsis thaliana*. Plant Biol. 10: 63–75.

Balazadeh, S., Siddiqui, H., Allu, D.A., Matallana-Ramirez, L., Caldana, C., Mehrnia, M. et al. (2010) A gene regulatory network controlled by the NAC transcription factor ANAC092/AtNAC2/ORE1 during salt-promoted senescence. Plant J. 62: 250–264.

Bohnert, J., Nelson, E., and Jensen, G. (1995) Adaptations to environmental stresses. Plant Cell. 7: 1099–1111.

Borisy, G.G. and Svitkina, T.M. (2000) Actin machinery: pushing the envelope. Curr. Opin. Cell Biol. 12: 104–112.

Chang, L., Azzolin, L., Biagio, D.D., Zanconato, F., Battilana, G., Xiccato, L.R. et al. (2018) The SWI/SNF complex is a mechanoregulated inhibitor of YAP and TAZ. Nature 563: 265–269.

Clement, M., Ketelaar, T., Rodiuc, N., Banora, M.Y., Smertenko, A., Engler, G. et al. (2009) Actin-depolymerizing factor2-mediated actin dynamics are essential for root-knot nematode infection of Arabidopsis. Plant Cell. 21: 2963–2979.

Cruz, J.R., and Moreno, Diaz de la Espina S. (2009) Subnuclear compartmentalization and function of actin and nuclear Myosin I in plants. Chromosoma. 118: 193–207

Dhar, N., Caruaba, J., Erdem, I., Subbarao, V.K., Klosterman, J.S., and Raina, R. (2020) The *Arabidopsis* SENESCENCE-ASSOCIATED GENE 13 Regulates Dark-Induced Senescence and Plays Contrasting Roles in Defense Against Bacterial and Fungal Pathogens. MPMI 33: 754–766.

Ding, S., Cheng, X., Wang, D., Chen, C., Yang, S., Wang, J. et al. (2022) *Aphelenchoides besseyi* Ab-FAR-1 Interacts with *Arabidopsis thaliana* AtADF3 to Interfere with Actin Cytoskeleton, and Promotes Nematode Parasitism and Pathogenicity. Int. Mol. Sci. 20: 12280.

Dingova, H., Fukalova, J., Maninova, M., Philimonenko, V.V., and Hozak, P. (2009) Ultrastructural localization of actin and actin-binding proteins in the nucleus. Histochem. Cell Biol. 131: 425–434.

Dong, H., Xia, X. and Hong, Y. (2001) ADF proteins are involved in the control of flowering and regulate F-actin organization, cell expansion, and organ growth in Arabidopsis. Plant Cell 13: 1333–1346.

Eckstein, A., Grzyb, J., Hermanowicz, P., Zglobicki, P., Labuz, J., Strzalka, W. et al. (2021) *Arabidopsis* Phototropins Participate in the Regulation of Dark-Induced Leaf Senescence. Int. J. Mol. Sci. 22: 1836.

Fernandez, K.M., Sinha, M., Zidan, M., and Renz, M. (2024) Nuclear actin filaments – a historical perspective. Nucleus 15: 2320656.

Guo, Y., Cai, Z., and Gan, S. (2004) Transcriptome of Arabidopsis leaf senescence. Plant Cell Environ. 27: 521–549.

Hörtensteiner, S., and Kräutler, B. (2011) Chlorophyll breakdown in higher plants. Biochim. Biophys. Acta. 1807: 977–988

Inada, N., Higaki, T., and Hasezawa, S. (2016) Nuclear function of subclass I Actin Depolymerizing Factor contributes to susceptibility in Arabidopsis to an adapted powdery mildew fungus. Plant Physiol. 170: 1420–1434.

Inada, N. (2017) Plant Actin Depolymerizing Factor – Actin microfilament disassembly and more. J. Plant Res. 130: 227–238.

Inada, N., Takahashi, N., and Umeda, M. (2021) Arabidopsis thaliana subclass I ACTIN DEPOLYMERIZING FACTORs and vegetative ACTIN2/8 are novel regulators of endoreplication. J. Plant Res. 134: 1291–1300.

Kandasamy, M.K., McKinney, E.C., and Meagher, R.B. (2009) A single vegetative actin isovariant overexpressed under the control of multiple regulatory sequences in sufficient for normal Arabidopsis development. Plant Cell 21: 701–718.

Kandasamy, M.K., McKinney, E.C., and Meagher, R.B. (2010) Differential sublocalization of actin variants within the nucleus. Cytoskeleton 67: 729–743.

Kim, Y.S., Sakuraba, Y., Han, S.H., Yoo, S.C., and Paek, N.C. (2013) Mutation of the Arabidopsis NAC016 Transcription Factor Delays Leaf Senescence. Plant Cell Physiol. 54: 1660–1672.

Kristó, I., Bajusz, I., Bajusz, C., Borkúti, P., and Vilmos, P. (2016) Actin, actin-binding proteins, and actin-related proteins in the nucleus. Histochem. Cell Biol. 145: 373–388.

Lichtenthaler, K. (1987) Chlorophylls and carotenoids: Pigments of photosynthetic biomembranes. Method. Enzymol. 148: 350–382.

Lim, O.P., Kim, J.H., and Nam, G.H. (2007) Leaf senescence. Annu. Rev. Plant Biol. 58: 115–136.

Lohman, K.N., Gan, S., John, M.C., and Amasino, R.M. (1994) Molecular analysis of natural leaf senescence in *Arabidopsis thaliana*. Physiol. Plant. 92: 322–328.

Matsumoto, T., Higaki, T., Takatsuka, H., Kutsuna, N., Ogata, Y., Hasezawa, S. et al. (2023) *Arabidopsis thaliana* subclass I ACTIN DEPOLYMERIZING FACTORs regulate nuclear structure and gene expression. Plant Cell Physiol. 10: 1231–1242.

Miao, Y., Laun, T., Zimmermann, P., and Zentgraf, U. (2004) Targets of the WRKY53 transcription factor and its role during leaf senescence in *Arabidopsis*. Plant Mol. Biol. 55: 853–867.

McDowell, J.M., Huang, S., McKinney, E.C., An, Y-Q., and Meagher, R.B. (1996a) Structure and evolution of the actin gene family in *Arabidopsis thaliana*. Genetics 142: 587–602.

McDowell, J.M., An, Y-Q., Huang, S., McKinney, E.C., and Meagher, R.B. (1996b) The Arabidopsis *ACT7* actin gene is expressed in rapidly developing tissues and responds to several external stimuli. Plant Physiol. 111: 699–711.

Meagher, R.B., McKinney, E.C., and Vitale, A.V. (1999) The evolution of new structures: clues from plant cytoskeletal genes. Trends Genet. 15: 278–284

Ojangu, E., Ilau, B., Tanner, K, Talts, K., Ihoma, E., Dolja, V. et al. (2018) Class XI Myosins Contribute to Auxin Response and Senescence-Induced Cell Death in *Arabidopsis*. Front. Plant Sci. 9: 1570.

Paavilainen, V.O., Bertling, E., Falck, S., and Lappalainen, P. (2004) Regulation of cytoskeletal dynamics by actin-monomer-binding proteins. Trends Cell Biol. 14: 386–394.

Peremyslov, V.V., Prokhnevsky, A.I., and Dolja, V.V. (2010) Class XI myosins are required for development, cell expansion, and F-actin organization in *Arabidopsis*. Plant Cell 22: 1883–1897.

Porter, K., Shimono, M., Tian, M., and Day, B. (2012) Arabidopsis Actin-Depolymerizing Factor-4 links pathogen perception, defense activation and transcription to cytoskeletal dynamics. PLoS Pathog. 8: e1003006.

Quirino, B.F., Normanly, J., and Amasino, R.M (1999) Diverse range of gene activity during *Arabidopsis thaliana* leaf senescence includes pathogen-independent induction of defense-related genes. Plant Mol. Biol. 40: 267–278.

Ren, G., An, K., Liao, Y., Zhou, X., Cao, Y., Zhao, H. et al. (2007) Identification of a Novel Chloroplast Protein AtNYE1 Regulating Chlorophyll Degradation during Leaf Senescence in Arabidopsis. Plant Physiol. 144: 1429–1441

Ruzicka, D.R., Kandasamy, M.K., McKinney, E.C., Burgos-Rivera, B., and Meagher, R.B. (2007) The ancient subclasses of Arabidopsis ACTIN DEPOLYMERIZING FACTOR genes exhibit novel and differential expression. Plant J. 52: 460–472.

Sakuraba, Y., Schelbert, S., Park, S.Y., Han, S.H., Lee, B.D., Andrès, C.B. et al. (2012) STAY-GREEN and chlorophyll catabolic enzymes interact at light-harvesting complex II for chlorophyll detoxification during leaf senescence in *Arabidopsis*. Plant Cell 24: 507–518.

Schelbert, S., Aubry, S., Burla, B., Agne, B., Kessler, F., Krupinska, K. et al. (2009) Pheophytin Pheophorbide Hydrolase (Pheophytinase) Is Involved in Chlorophyll Breakdown during Leaf Senescence in *Arabidopsis*. Plant Cell 21: 767–785.

Shimoda, Y., Ito, H., and Tanaka, A. (2016) Arabidopsis *STAY-GREEN*, Mendel’s Green Cotyledon Gene, Encodes Magnesium-Dechelatase. Plant Cell 28: 2147–2160.

Tan, S., Sha, Y., Sun, L., and Li, Z. (2023) Abiotic Stress-Induced Leaf Senescence: Regulatory Mechanisms and Application. Int. J. Mol. Sci. 24: 11996.

Takatsuka, H., Higaki, T., and Umeda, M. (2018) Actin reorganization triggers rapid cell elongation in roots. Plant Physiol. 178: 1130–1141.

Tian, M., Chaudhry, F., Ruzicka, D.R., Meagher, R.B., Staiger, C.J., and Day, B. (2009) Arabidopsis Actin-Depolymerizing Factor AtADF4 mediates defense signal transduction triggered by the *Pseudomonas syringae* effector AvrPphB. Plant Physiol. 150: 815–824.

Wang, L., Qiu, T., Yue, J., Guo, N., He, Y., Han, X. et al. (2021) *Arabidopsis* ADF1 is Regulated by MYB73 and is Involved in Response to Salt Stress Affecting Actin Filament Organization. Plant Cell Physiol. 62: 1387–1395.

Wang, L., Cheng, J., Bi, S., Wang, J., Cgeng, X., Liu, S. et al. (2023) Actin Depolymerization Factor ADF1 Regulated by MYB30 Plays an Important Role in Plant Thermal Adaptation. Int. J. Mol. Sci. 24: 5675.

Weaver, L.M., Gan, S., Quirino, B., and Amasino, R.M. (1998) A comparison of the expression patterns of several senescence-associated genes in response to stress and hormone treatment. Plant Mol. Biol. 37: 455–469.

Weaver L.M. and Amasino R.M. (2001) Senescence Is Induced in Individually Darkened Arabidopsis Leaves, but Inhibited in Whole Darkened Plants. Plant Physiol. 127: 876–886.

Yao, H., Li, X., Peng, L., Hua, X., Zhang, Q., Li, K. et al. (2022) Binding of 14-3-3κ to ADF4 is involved in the regulation of hypocotyl growth and response to osmotic stress in Arabidopsis. Plant Sci. 320: 111261.

Zhang, Y., Liu, Z., Wang, X., Wang, J., Fan, K., Li, Z. et al. (2018) DELLA proteins negatively regulate dark-induced senescence and chlorophyll degradation in *Arabidopsis* through interaction with the transcription factor WRKY6. Plant Cell Rep. 37: 981–992.

Zhang, Y., Guo, P., Xia, X., Guo, H., and Li, Z. (2021) Multiple Layers of Regulation on Leaf Senescence: New Advances and Perspectives. Front. Plant Sci. 12: 788996.

