## Supplemental Materials for "*Arabidopsis thaliana ACTIN DEPOLYMERIZING FACTOR*s play a role in leaf senescence regulation"

**SUPPOTING INFORMATION**

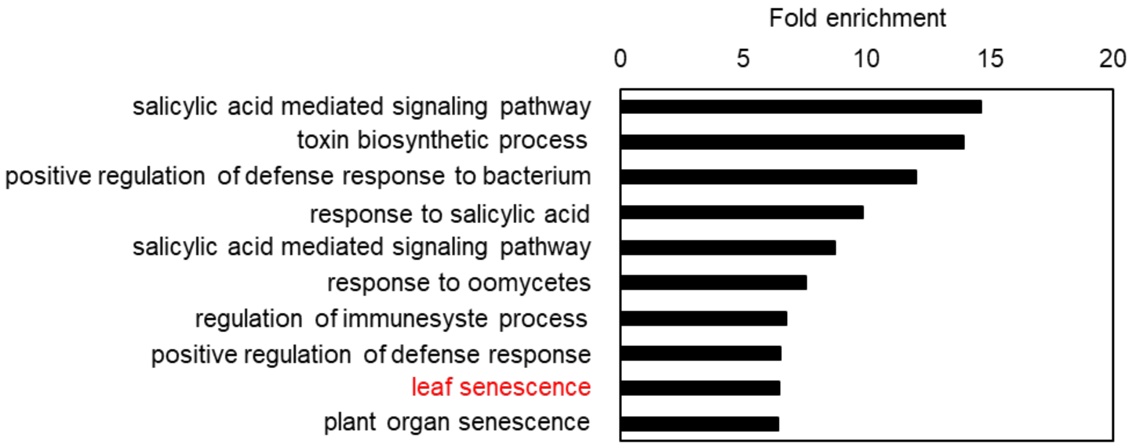


**Figure S1. The expression of leaf senescence genes was upregulated in *ADF1-4Ri*.** Enrichment analysis of GO terms for upregulated genes in *ADF1-4Ri* mature leaves.

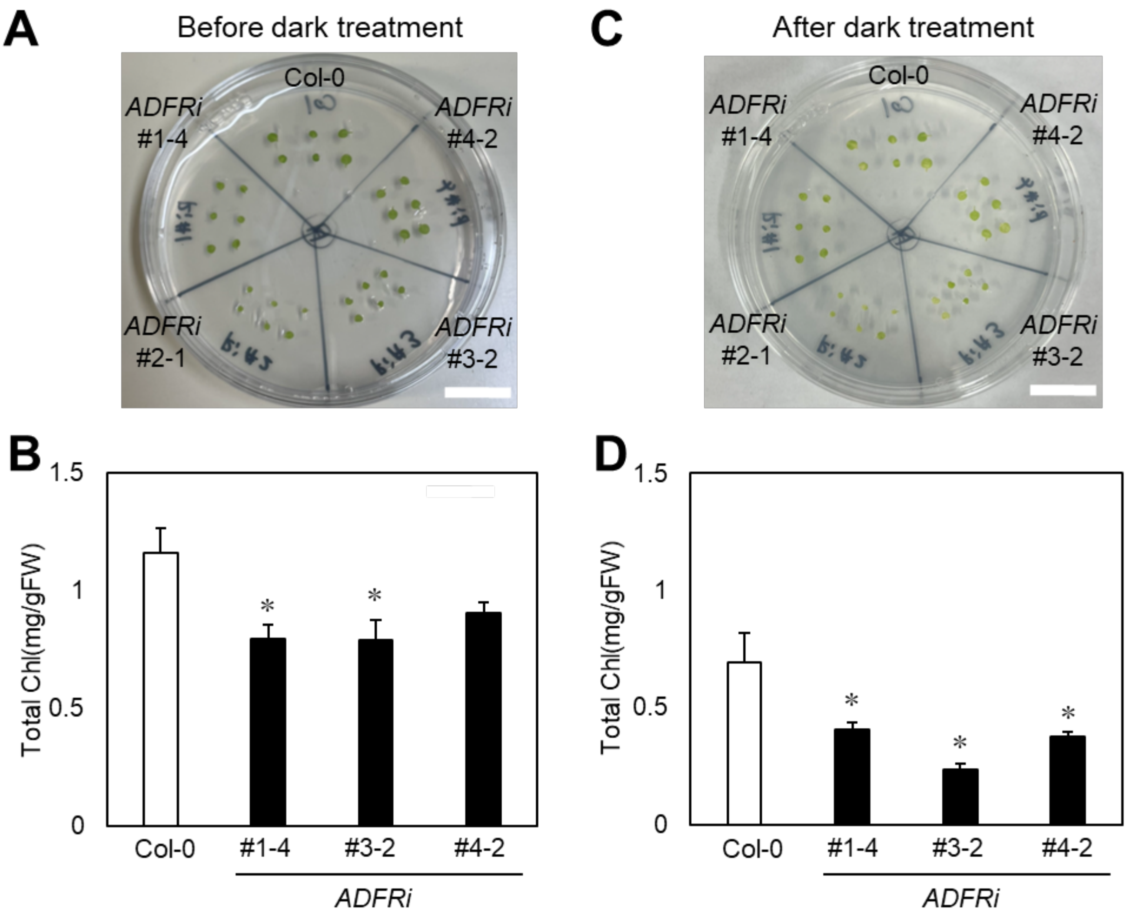


**Figure S2. Chlorophyll content was reduced in *ADF1-4Ri* lines before and after dark treatment.** (**A**) The first and second leaves excised from 21-day-old Col-0 and *ADF1-4Ri* (#1-4, #2-1, #3-2, and #4-1) plants grown at 23°C under short-day conditions. (**B**) Total chlorophyll contents of the first and second leaves of 21-day-old plants. (**C**) The first and second leaves were treated in the dark for 4 days. (**D**) Total chlorophyll contents of 21-day-old plants treated in the dark for 4 days. Bars in **A** and **C** indicate 1 cm. In **B** and **D**, the average values of three independent experiments were shown. Error bars indicate standard deviation (SD). Comparison with Col-0 was performed by Student's *t*-test. Asterisks indicate that there is a significant difference (* $P$<0.05).

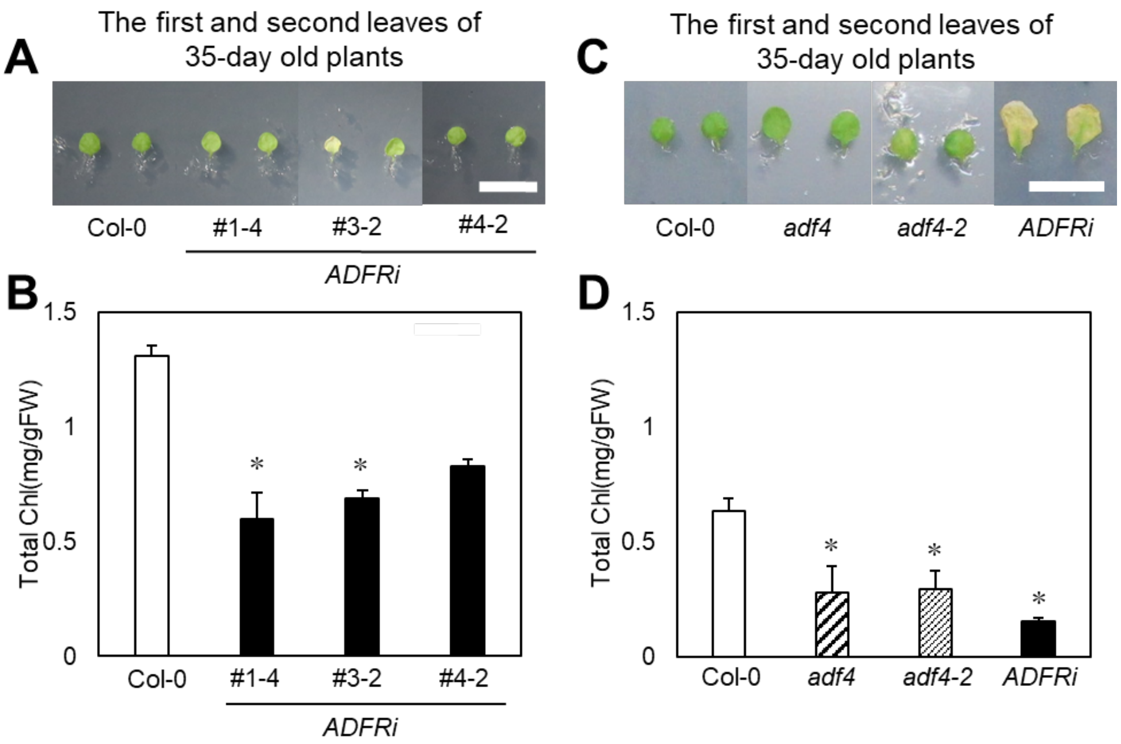


**Figure S3. *ADF1-4Ri* lines and *adf4* show early age-dependent leaf senescence.** (**A**) The first and the second leaves of 35-day-old Col-0 and *ADF1-4Ri* lines (#1-4, #3-2, and #4-1) grown at 23°C under short-day conditions. (**B**) Total chlorophyll contents of 35-day-old plants. (**C**) The first and the second leaves of 35-day-old Col-0, *adf4-1*, *adf4-2*, and *ADF1-4Ri*#2-1 grown at 23°C under short-day conditions. (**D**) Total Chlorophyll contents of the first and the second leaves of 35-day-old plants. Bars in **A** and **C** indicate 1 cm. For **B** and **D**, 3 independent experiments were performed, and the average values were shown. Error bars indicate SD. Comparison with Col-0 was performed by Student's *t*-test. Asterisks indicate that there is a significant difference (* $P<0.05$).

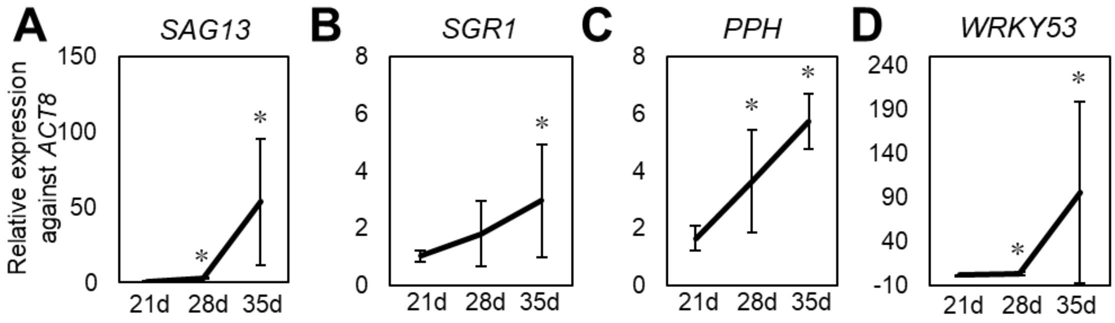


**Figure S4. The expression of *SAG*s was upregulated in Col-0 during age-dependent leaf senescence.** The expression of *SAG13* (**A**), *SGR1* (**B**), *PPH* (**C**), and *WRKY53* (**D**) in the first and the second leaves of plants at 21-, 28- and 35-days. For normalization, *ACT8* (At1g49290) was used as the internal control. The average values were obtained from more than two biological replicates, one biological replicate containing three technical replicates. Error bars indicate SD. Comparison with expression at 21-days was performed by Student's *t*-test. Asterisks indicate $P<0.05$.

**Table S1.** Primers for qRT-PCR.

| | | |
|---|---|---|
| ACT8 | For | TCAGCACTTTCCAGCAGATG |
| | Rev | ATGCCTGGACCTGCTTCAT |
| UBQ11 | For | ACCAGCAGCGTCTCATCTTC |
| | Rev | TGTAGTCGGCCAAAGTACGTC |
| SAG13 | For | GCTTTCCATCTCTCACAGCTTGCC |
| | Rev | ATTGACATGCACGACTCCAGC |
| SGR1 | For | TGGGCAAATAGGCTATACCG |
| | Rev | CCACCGCTTATGTGACAATG |
| PPH | For | TAAGGGTGTTTGGAGGGAAG |
| | Rev | AGGAGTTCCTTCGATTTAGGTG |
| WRKY53 | For | CGGAAGTCCGAGAAGTGAAG |
| | Rev | ATCTTGGCGATGATGACTCTCG |
| ADF1 | For | TGAGTGATGGTACTGGTACTTGA |
| | Rev | CAAGACCGAAACACCGATAGAATG |
| ADF2 | For | TCTGGAGTGAATATGTTTCCTCTG |
| | Rev | ACATACAATAATACCAAGTAGAATG |
| ADF3 | For | TCGGTTGAATCAAACTTTTTCGT |
| | Rev | GGTACCGTCACAGCAAACTTTAGG |
| ADF4 | For | GATACACTTGACACCCTTCATTCTATCT |
| | Rev | AGGAAGCAAACACAGCACAAACCTTGTG |

### SUPPLEMENTAL METHODS

#### Gene ontology enrichment

GO analysis was performed using PANTHER 17.0 (Released 20221013). The microarray data were published in Matsumoto et al., 2023, and deposited in the Gene Expression Omnibus (GEO) of National Center for Biotechnology Information (NCBI), accessible under the GEO accession number GSE228396.
